# CD4 T cell surface architecture is associated with response to anti-PD-1 therapy in non-small cell lung cancer

**DOI:** 10.64898/2026.10.26.761814

**Authors:** Riccardo Cinotti, Ying Yang, Vincent van Hoef, Hanna van Ooijen, Andreas Parling, Mannon Geindreau, Neda Bigdeli, Mireia Cruz de Los Santos, Lucas Baldran-Groves, Yiming Yang, Lisa Liu, Maria Johansson, Klas Kärre, Simon Fredriksson, Luigi de Petris, Andreas Lundqvist

## Abstract

Anti-PD-1 therapy has substantially improved outcomes for patients with advanced non-small cell lung cancer (NSCLC), but clinical responses remain highly heterogeneous and are difficult to predict. Current immune profiling approaches primarily characterize immune cell composition and protein abundance, whereas the spatial organization of proteins at the cell surface has remained largely unexplored. Here, we demonstrate for the first time that cell surface protein organization is associated with clinical response to immune checkpoint inhibition therapy. Using the Proximity Network Assay (PNA), we simultaneously quantified cell frequencies, protein abundance, and protein organization in pretreatment peripheral blood mononuclear cells from 62 patients with advanced NSCLC treated with anti-PD-1 agents. While most cell populations were unchanged between responders and non-responders, pronounced response-associated alterations were identified within the CD4 T-cell compartment. Responders exhibited elevated expression of adhesion-associated proteins, and remodeling of a CD5-containing membrane domain. Increased colocalization within this domain was associated with poor clinical outcome and reduced survival. Machine learning analysis further identified CD4 T cell features as the strongest predictors of treatment response and demonstrated that protein organization provided greater predictive power than protein abundance alone. Together, these findings identify cell surface protein organization as a previously unexplored source of biomarkers and highlight its potential to improve immune profiling and biomarker discovery in cancer immunotherapy.

## Introduction

Immune checkpoint inhibition targeting the programmed cell death protein 1 (PD-1) pathway has transformed the treatment landscape for multiple malignancies, including non-small cell lung cancer (NSCLC)^1–3^. However, despite producing durable clinical responses in a subset of patients, a large proportion of individuals fail to benefit from treatment, highlighting the need for improved biomarkers that can identify responders and provide insight into the mechanisms underlying therapeutic success and resistance^4–6^.

Current biomarkers for anti-PD-1 therapy, including tumor PD-L1 expression, tumor mutational burden, and immune cell infiltration, provide only partial predictive value^7–9^. Due to its ease of sampling and that anti-tumor immunity emerges from coordinated interactions between multiple immune cell populations, increasing attention has been directed towards the systemic immune compartment as a source of clinically relevant biomarkers. Peripheral blood offers a minimally invasive window into immune function and has previously revealed associations between treatment outcome and the abundance and activation status of circulating lymphoid and myeloid populations^10–13^. Most of these studies have focused on cellular composition or RNA and protein abundance, but higher-order features of immune cell phenotype remain largely unexplored.

The function of immune cells is not solely determined by which proteins they express, but also by how these proteins are organized within the plasma membrane^14^. Spatial organization regulates receptor signaling, cell-cell interactions, migration, and immune synapse formation through the assembly of molecular complexes and membrane domains^15–18^. Alterations in receptor colocalization and membrane architecture have been shown to influence immune cell activation and effector function, suggesting that spatial protein organization may provide complementary biological information beyond conventional expression measurements^19,20^.

To investigate the relationship between peripheral immune cell surface architecture and clinical response to anti-PD-1 therapy, we applied the Proximity Network Assay (PNA) to Peripheral Blood Mononuclear Cells (PBMCs) collected from patients with NSCLC prior to treatment initiation. This approach simultaneously quantifies immune cell composition, protein abundance, and protein organization, at single cell resolution^21^. By integrating these complementary layers of information across major immune cell populations, we sought to describe the membrane architecture of peripheral immune cells, identify cellular and molecular features associated with therapeutic response, and to evaluate the potential of membrane protein organization as a novel source of immunotherapy biomarkers.

## Results

### Clinical response to anti-PD-1 therapy is associated with CD8 T cell pretreatment frequencies

To identify features of peripheral immune cells associated with response to PD-1 blockade, pretreatment PBMC samples from 62 patients with advanced NSCLC receiving anti-PD-1 therapy were analyzed by PNA (Figure 1A, Table 1). Patients were classified according to RECIST criteria as objective responders (OR; complete or partial response, n = 29) or non-responders (non-OR; stable or progressive disease, n = 33). Responders exhibited significantly improved overall survival (OS) compared with non-responders (Figure 1B), and analysis of individual RECIST categories demonstrated a stepwise relationship between response and survival (Figure 1C). Age, sex, chemotherapy and PD-L1 status did not significantly associate with response (Figure S1A).

**Figure 1.**
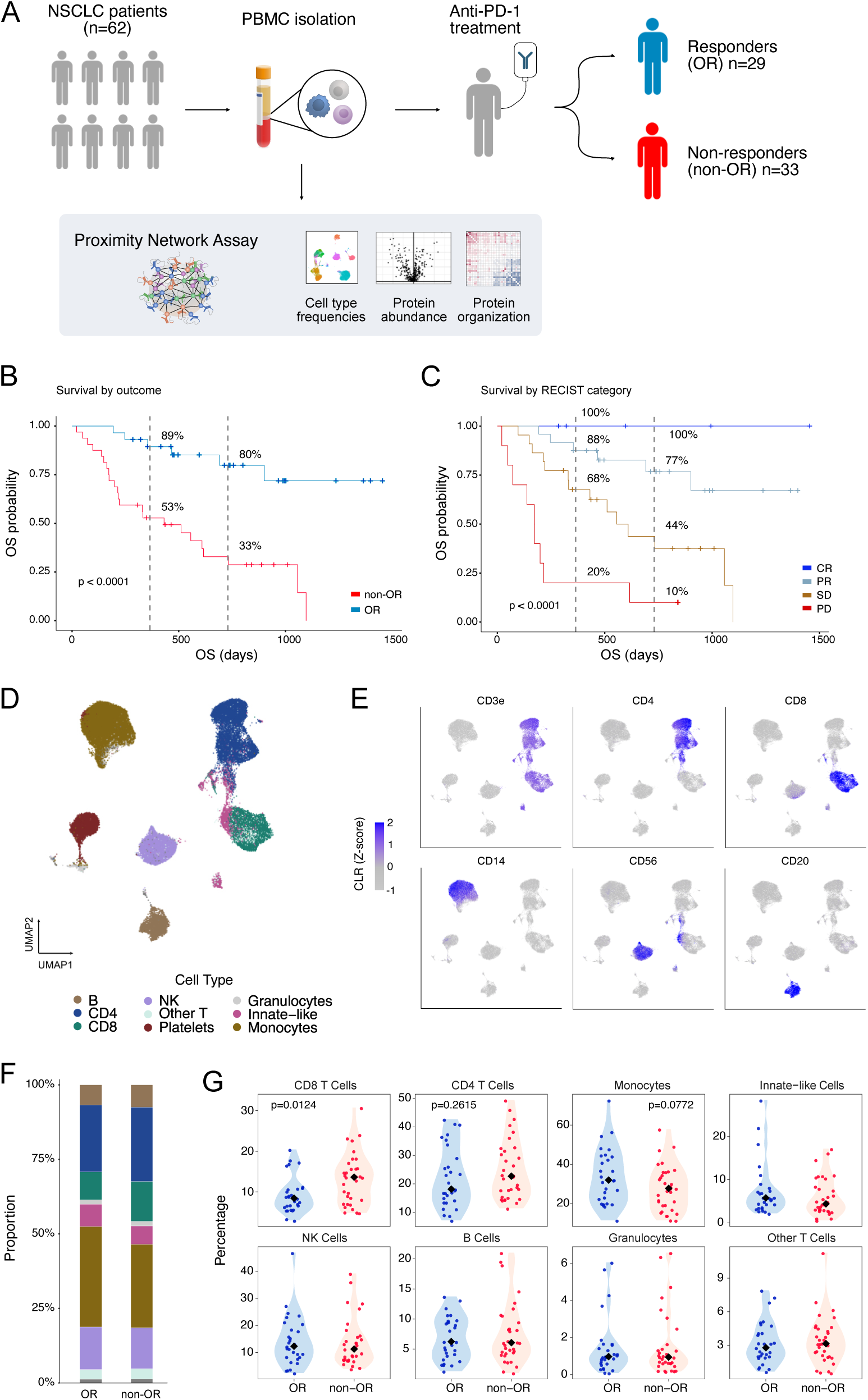
Study Overview. (**A**) Schematic representation of the experimental design and workflow. PBMCs from 29 objective responders (OR) and 33 non-responders (non-OR) were collected prior to anti-PD-1 therapy and analyzed using the Proximity Network Assay (PNA). (**B**) Kaplan–Meier survival analysis comparing overall survival between OR and non-OR; 1- and 2-year survival rates are indicated by vertical dashed lines. (**C**) Kaplan–Meier survival curves stratified by RECIST clinical response criteria; 1- and 2-year survival rates are indicated by dashed lines. (**D**) UMAP visualization of the entire PBMC dataset, highlighting major immune cell compartments. (**E**) Relative abundance of lineage-defining proteins across the full dataset. (**F**) Proportion of major immune cell populations, stratified by patient response group. (**G**) Distribution of cell type proportions per patient, categorized by response status. Statistical significance was evaluated using a two-tailed t-test. Each dot represents a single patient and p-values < 0.1 are indicated. Other T cells are CD3e positive cells that could not be assigned to any of the conventional T cell subpopulations together with double positive T cells. The innate-like lymphocyte compartment is composed of gamma-delta T cells, MAIT, NKT-like cells and double negative T cells.

**Table 1.** Clinical metadata.

| Clinical metadata |  | OR (n=29) | non-OR (n=33) |
| --- | --- | --- | --- |
| Age, median (range), yr |  | 70 (51-84) | 69 (32-78) |
| Sex, n (%) |  |  |  |
|  | Male | 12 (41%) | 17 (52%) |
|  | Female | 17 (59%) | 16 (48%) |
| Smoking, n (%) |  |  |  |
|  | Current | 6 (21%) | 7 (21%) |
|  | Former | 15 (52%) | 24 (73%) |
|  | Non-smoker | 8 (28%) | 2 (6%) |
| Histologic type, n (%) |  |  |  |
|  | Adenocarcinoma | 29 (100%) | 33 (100%) |
| PD-L1 expression, n (%) |  |  |  |
|  | Not analyzed | 1 (3%) | 1 (3%) |
|  | < 1% | 7 (24%) | 19 (58%) |
|  | 1-49% | 8 (28%) | 5 (15%) |
|  | >50% | 13 (45%) | 8 (24%) |
| Therapy, n (%) |  |  |  |
|  | Anti-PD1 | 8 (28%) | 5 (15%) |
|  | Anti-PD1 + chemotherapy | 21 (72%) | 27 (85%) |
| Type of ICI, n (%) |  |  |  |
|  | Pembrolizumab | 20 (69%) | 24 (73%) |
|  | Cemiplimab | 9 (31%) | 9 (27%) |
| Reason for treatment stop, n (%) |  |  |  |
|  | Ongoing | 3 (10%) | 3 (9%) |
|  | Planned | 3 (10%) | - |
|  | Progression | 13 (45%) | 20 (61%) |
|  | Toxicity | 10 (34%) | 10 (30%) |
| Best overall response, n (%) |  |  |  |
|  | CR | 5 (17%) | - |
|  | PR | 24 (83%) | - |
|  | SD | - | 23 (70%) |
|  | PD | - | 10 (30%) |
| Progression-free survival (PFS) |  |  |  |
|  | PFS, median days (95%CI) | 616 (351- ) | 172 (97-302) |
|  | PFS, event observed, n (%) | 17 (59%) | 30 (91%) |
| Overall survival (OS) |  |  |  |
|  | OS, median days (95% CI) | Not reached | 511 (214-733) |
|  | OS, event observed, n (%) | 9 (31%) | 25 (76%) |

High-dimensional clustering and cell annotation based on canonical surface markers identified all major PBMC populations, including CD4 T cells, CD8 T cells, NK cells, B cells, monocytes, granulocytes, and smaller innate immune populations (Figure 1D-E and Figure S1B). Although immune cell composition was largely similar between responders and non-responders, responders exhibited significantly lower frequencies of CD8 T cells together with trends towards increased monocyte frequencies and reduced CD4 T cell frequencies (Figure 1F-G). These findings motivated a more detailed characterization of the major immune cell compartments.

### CD8 T cells exhibit conserved phenotypic and organizational profiles irrespective of treatment outcome

Given the reduced frequencies of circulating CD8 T cells in responders (Figure 1G), an in-depth characterization of the CD8 T cell compartment was performed. Re-clustering combined with conventional gating of CD8 T cells identified six distinct cell subsets corresponding to canonical differentiation states, including Naïve, Central Memory (TCM), Effector Memory (TEM), and Effector Memory Re-expressing CD45RA (TEMRA) populations (Figure 2A-B, Figure S1B). These subsets displayed expected expression patterns of costimulatory and differentiation markers, with progressive loss of CD127 and CD28 and increased expression of Fas, KLRG1, and CD57 along the differentiation trajectory from naïve to terminally differentiated cells (Figure 2C-D). Additionally, PD-1 expression was highest within memory CD8 populations, particularly TCM cells. However, despite the reduction in overall CD8 T cell frequency, no significant differences were observed between responders and non-responders in the relative frequency of the individual T cell subsets or their phenotypes (Figure S2A-C).

**Figure 2.**
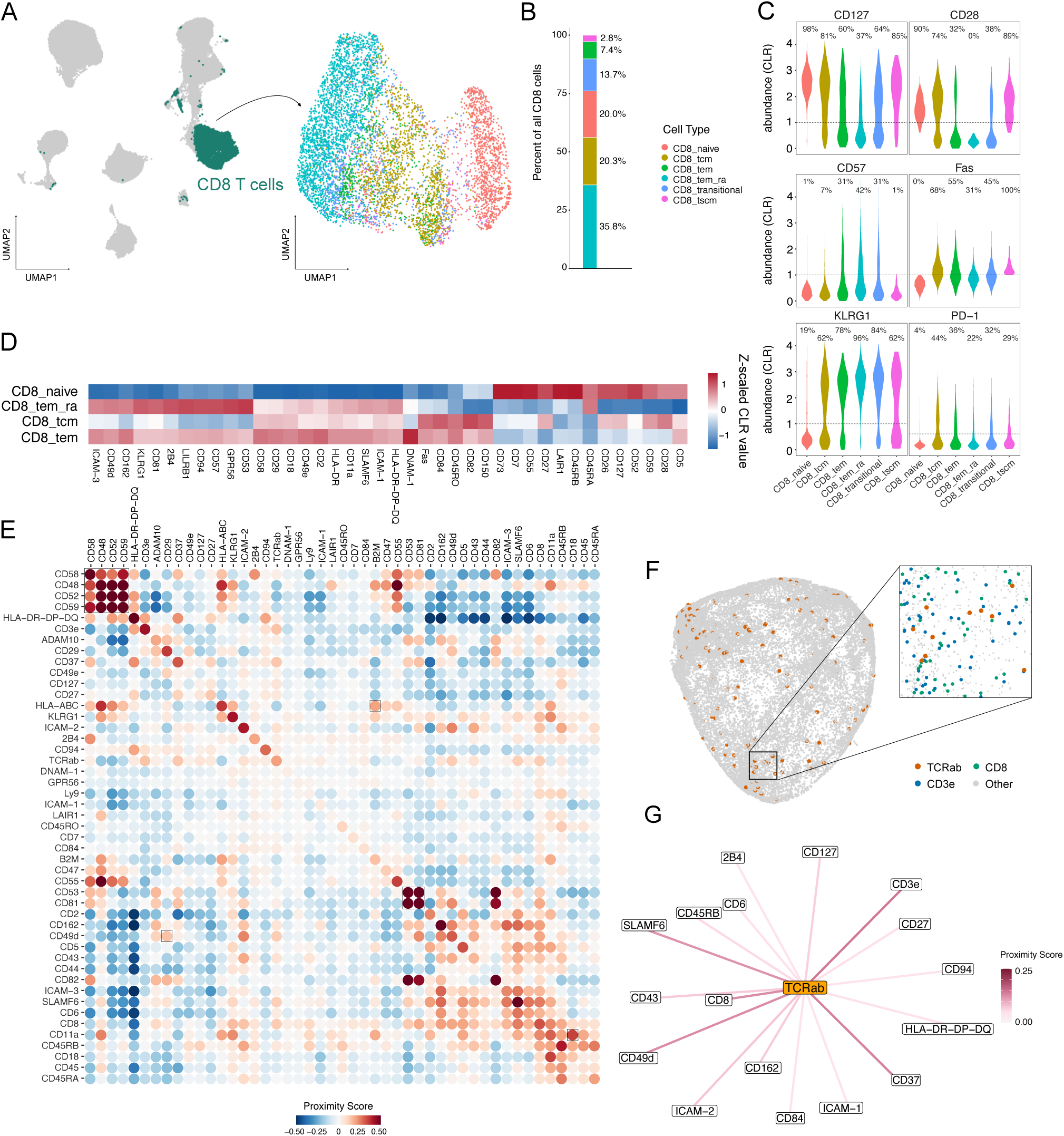
CD8 T cell characterization. (**A**) UMAP visualization of the re-clustered CD8 T cell compartment, identifying distinct cellular subpopulations. (**B**) Percentage of CD8 T cell subpopulations within the total CD8 compartment. (**C**) Protein abundance distribution across CD8 T cell subpopulations; percentage of cells above the threshold are indicated. (**D**) Heatmap showing protein abundance, expressed as Z-scaled centered log-ratio (CLR) values, for the most variable proteins across Naive, TCM, TEM and TEMRA populations. (**E**) Median per-patient mean proximity score (log2 ratio) for relevant CD8 T cell proteins. A selection of well-known complexes is highlighted. (**F**) Representative visualization of TCRαβ on a single CD8 T cell (left) and of CD3e and CD8 in a TCRαβ-rich neighborhood on this cell (right). (**G**) Positive colocalization network of TCRαβ, showing median proximity scores across patient samples.

Next, the membrane organization within the CD8 compartment was characterized. This analysis identified several well-established membrane complexes and domains, including LFA-1 (CD11a/CD18), VLA-4 (CD49d/CD29), B2M-HLA-ABC, and a cluster of GPI-anchored proteins known to associate with lipid rafts (Figure 2E). Focusing on the TCRαβ protein, network analysis revealed known interaction partners including CD3ε and CD8 (Figure 2E-F). In addition, PNA identified several additional known and novel interaction partners of the TCR, including adhesion proteins CD49d and CD43 and regulatory molecules CD6 and 2B4 (Figure 2G), demonstrating the ability of PNA to resolve membrane protein organization in primary immune cells. Overall, although demonstrating a strongly defined membrane structure, the CD8 T cell spatial phenotype was not different between responders and non-responders (Figure S2D-E). These results suggest that, although the frequency of CD8 T cells correlates with outcome, most of their expression profile and membrane architecture is preserved irrespective of clinical outcome.

### Clinical response to PD-1 therapy is associated with CD5 remodeling in circulating CD4 T cells

In addition to the same six distinct T cell subsets identified for CD8 T cells, regulatory T cells (Tregs) were identified upon re-clustering of CD4 T cells (Figure 3A-B). All the populations displayed expected differentiation-associated marker profiles (Figure 3C-D), and within CD4 subsets, PD-1 was primarily expressed by effector memory CD4 T cells. Responders exhibited a trend towards enrichment of effector memory CD4 T cells, together with increased TCM cells and reduced naïve cells, suggesting a shift towards a more differentiated CD4 T cell compartment (Figure S3A).

**Figure 3.**
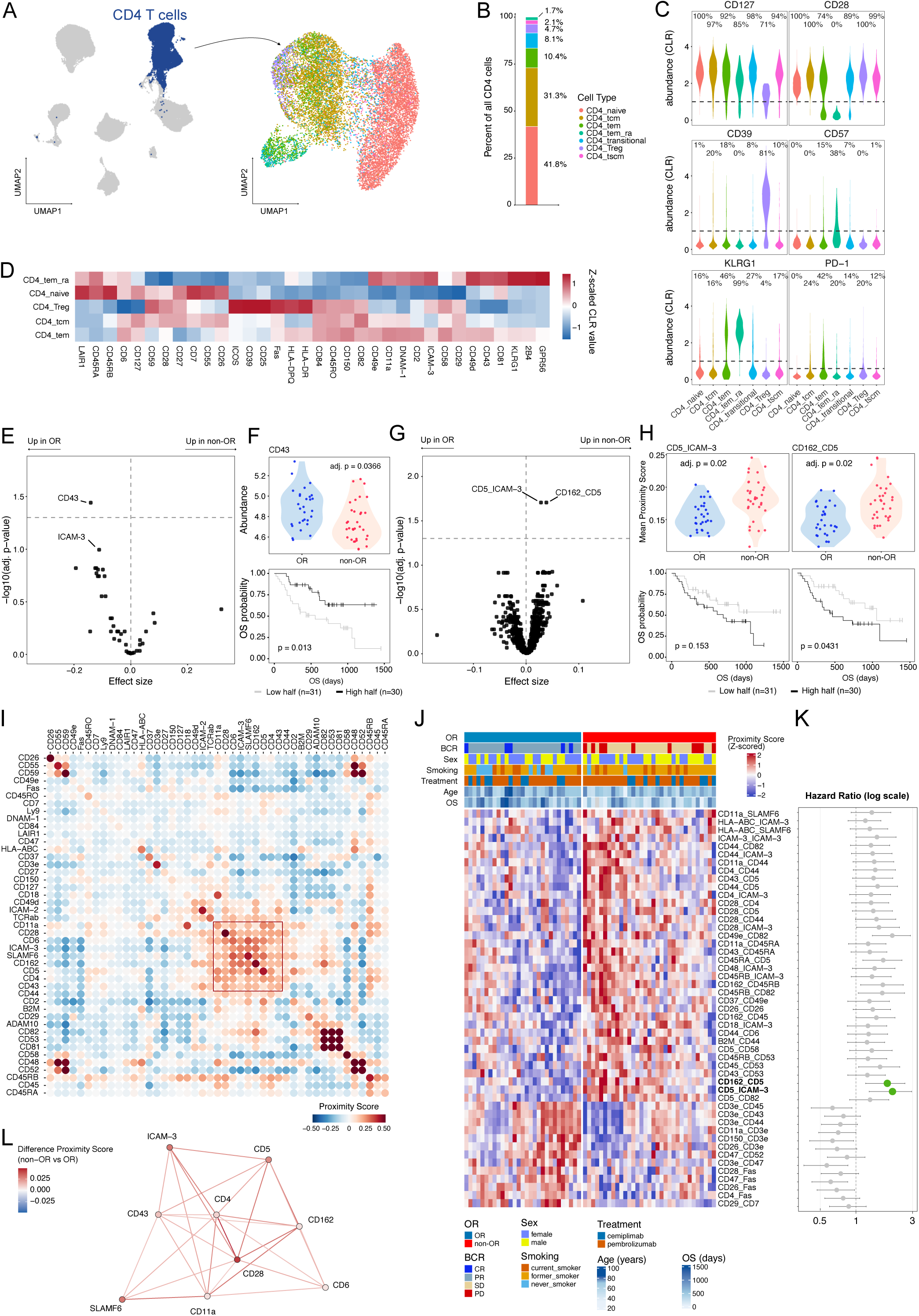
Profiling and clinical association of the CD4 T cell compartment. (**A**) UMAP projection of re-clustered CD4 T cells, identifying distinct cellular subpopulations. (**B**) Relative frequency of identified CD4 T cell subpopulations within the total CD4 compartment. (**C**) Distribution of protein expression levels across CD4 T cell subpopulations; percentages exceeding the detection threshold are indicated. (**D**) Heatmap of protein abundance, expressed as Z-scaled centered log-ratio (CLR) values, for the most variable proteins across Naive, TCM, TEM, TEMRA and Treg subpopulations. (**E**) Volcano plot showing differentially abundant proteins between OR and non-OR; horizontal line denotes adjusted p-value < 0.05. (**F**) Abundance distribution of significant marker with each dot representing the mean CLR of a patient (top) and Kaplan–Meier survival analysis comparing patients stratified by the highest versus lowest half of marker abundance (bottom). (**G**) Volcano plot highlighting differentially colocalized marker pairs between OR and non-OR patients; horizontal line denotes adjusted p-value < 0.05. (**H**) Proximity Score distribution of significant markers with each dot representing the mean log2 ratio of a patient (top) and Kaplan–Meier survival analysis comparing patients stratified by the highest versus lowest half of mean log2 ratio (bottom). (**I**) Median per-patient mean proximity score (log2 ratio) for relevant CD4 T cell markers. The box highlights a large protein domain containing CD5. (**J**) Z-scaled proximity scores for significantly differentially colocalized marker pairs (OR vs non-OR, p < 0.01). (**K**) Cox proportional hazards forest plot illustrates the hazard ratio (95% CI) for overall survival per 1 standard deviation increase in mean log2 ratio for the marker pairs presented in J; dashed line denotes HR = 1 and green dots the significant marker pairs in the OR versus non-OR comparison from panel G (right). (**L**) Network visualization of the differential marker proximity for markers highlighted in (**I**). Nodes and edges are colored according to the effect size of the non-OR vs OR difference in marker-pair proximity.

Analysis of the correlation between CD4 T cell protein abundance and clinical response revealed that the expression of several adhesion proteins was related to outcome. Within the whole CD4 T cell compartment, CD43 was significantly higher in responders, and ICAM-3 showed trends toward increased abundance compared with non-responders (Figure 3E-F, Figure S3B). Notably, CD49e and CD47 were significantly upregulated in responder CD4 TEM and TEMRA cells respectively (Figure S3C). Higher CD43 and ICAM-3 abundance was also associated with improved overall survival, supporting their association with clinical benefit (Figure 3F, Figure S3B).

Given these coordinated phenotypic changes, cell surface architecture was subsequently examined (Figure 3G-L). Differential proximity analysis revealed two co-localizations, CD5–ICAM-3 and CD5–CD162, to be significantly increased in non-responders (Figure 3G-H, Figure S3D, Supplementary video 1). This phenotype was particularly evident in naive CD4 T cells (Figure S3E). Furthermore, network analysis revealed a highly interconnected protein domain centered around CD5 and enriched for adhesion molecules and regulators of T cell activation (Figure 3I). Although only two protein pairs displayed significant alterations after multiple-testing correction, a substantially larger set of interactions showed coordinated variation across donors with non-adjusted p-values < 0.01 (Figure 3J). Many of these alterations involved the same group of proteins, part of the identified protein domain, including CD5, ICAM-3, CD162, CD11a, CD43, CD44, CD6, CD28 and SLAMF6, suggesting remodeling of a broader membrane structure rather than isolated pairwise changes. Notably, increased colocalization was seen for all proteins within this domain in non-responding patients (Figure 3L), and the alteration was indeed associated with reduced overall survival (Figure 3K). Together, these findings identify a response-associated CD4 T cell state characterized by increased memory differentiation, altered adhesion protein expression and remodeling of a CD5-centered membrane domain.

### Clinical response to anti-PD-1 therapy is associated with altered monocyte composition and platelet-monocyte interactions

Next, the monocyte compartment was characterized to determine whether response-associated features were present within the myeloid lineage. Three canonical monocyte subsets were identified, comprising classical, intermediate, and non-classical monocytes, with classical monocytes being the dominant population (Figure 4A-B). Comparison of subset frequencies revealed a significant enrichment of non-classical monocytes in responders (Figure 4C). No significant differences were observed for classical or intermediate monocytes (Figure S4A).

**Figure 4.**
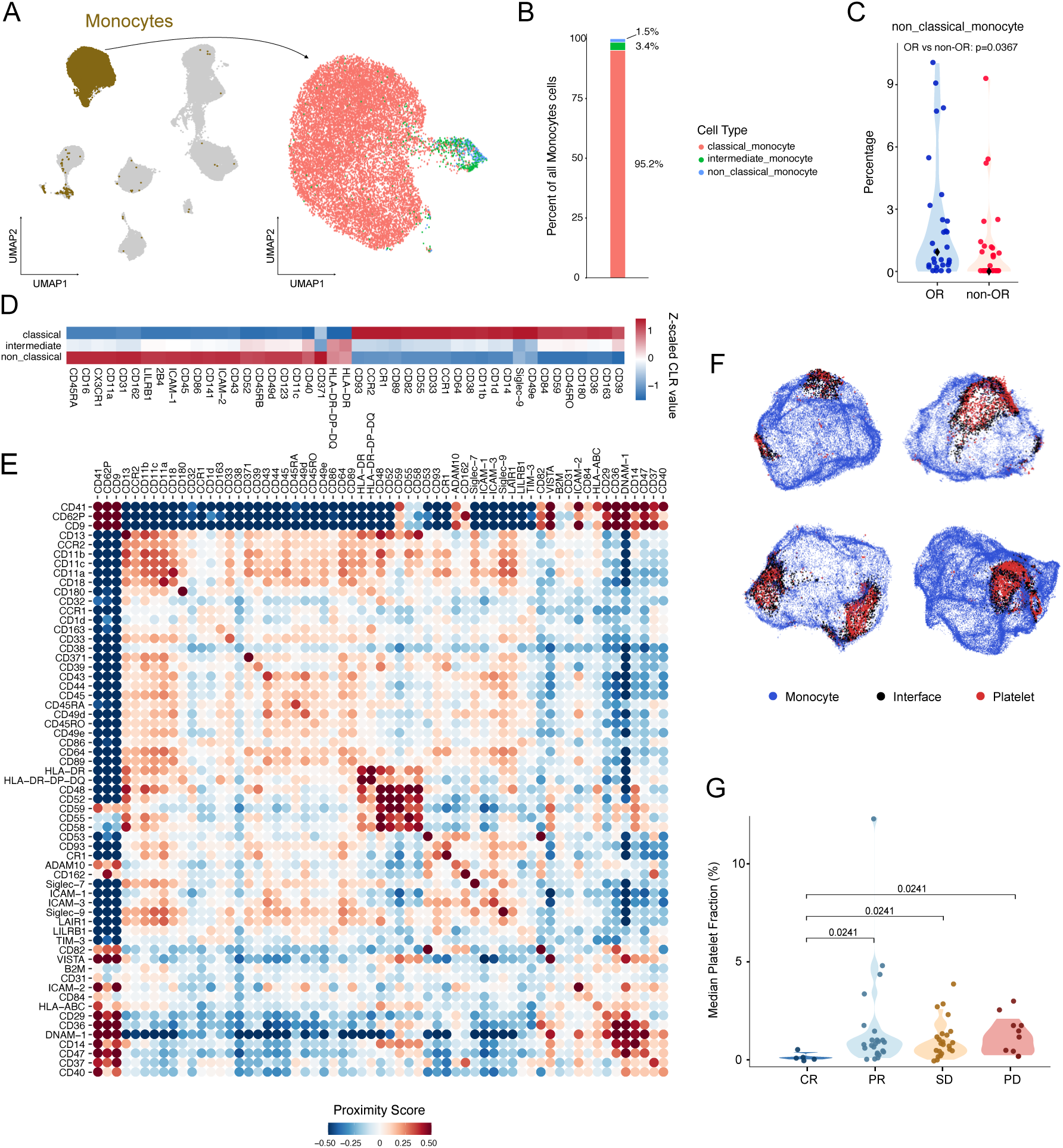
Profiling of the monocyte compartment. (**A**) UMAP projection of re-clustered monocytes, identifying distinct cellular subpopulations. (**B**) Relative frequency of monocyte subpopulations within the total monocyte compartment. (**C**) Quantification of non-classical monocyte proportions within the total monocyte compartment in OR vs non-OR patients. Each dot represents the percentage of a single patient (**D**) Heatmap of protein abundance, expressed as Z-scaled centered log-ratio (CLR) values, for the most variable proteins across monocyte subpopulations. (**E**) Median per-patient mean proximity score (log2 ratio) for relevant monocyte markers. (**F**) Representative visualization of platelet-monocyte interactions, highlighting platelets (red) on four monocyte cells (blue). (**G**) Quantification of the fraction of nodes assigned to platelets per RECIST clinical response category. Individual dots indicate the median fraction per patient (based on a subsample of 1000 random monocytes).

Although all three subsets displayed distinct protein expression profiles consistent with their established phenotypes (Figure 4D), neither protein abundance nor protein organization analyses identified significant response-associated differences (Figure S4B-E). However, analysis of monocyte membrane organization revealed a distinct cluster consisting of the platelet markers CD41, CD62P and CD9 (Figure 4E). The segregation of these markers from the broader monocyte network indicated the presence of platelet-monocyte aggregates (PMAs), which was further supported by visualization of individual monocytes containing platelet-associated protein nodes (Figure 4F).

Although not significantly different between responders and non-responders (Figure S4F), quantification of PMAs demonstrated differences across RECIST response groups, with complete responders exhibiting the lowest levels and significantly fewer PMAs than patients with progressive disease (Figure 4G). Notably, PMAs were almost exclusively observed within the monocyte compartment and were largely absent from T cells, NK cells, and B cells (Figure S4G).

### NSCLC patients display a heterogeneous NK compartment that does not predict clinical response

Finally, the NK cell compartment was investigated. Although NK cells are conventionally classified according to CD56 and CD16 expression into CD56^bright^ CD16^-^ and CD56^dim^ CD16^+^ cells, overlaying these canonical cell populations onto the PNA-derived UMAP showed that the defined cell types resulted in poor separation and that additional subpopulations were overlooked (Figure S5A-D). Therefore, unsupervised clustering was carried out, identifying seven distinct NK cell populations (Figure 5A-B), including CD56^bright^ cells (cluster 5), memory-like NKG2C^+^ NK cells (cluster 4), differentiated CD57^+^ NK cells (cluster 6), and additional phenotypically distinct subsets (clusters 0-3) defined by coordinated differences in activating receptors, inhibitory receptors, adhesion molecules, and differentiation markers (Figure 5C-D). These results indicate that the PNA analysis captured NK cell heterogeneity at greater depth than conventional phenotyping, and the identified phenotypes correlate well with previously reported NK cell subgroups^22^. Despite this phenotypic diversity, no significant differences were detected between responders and non-responders in NK cell subset frequencies, protein abundance, or protein organization (Figure S5E-I).

**Figure 5.**
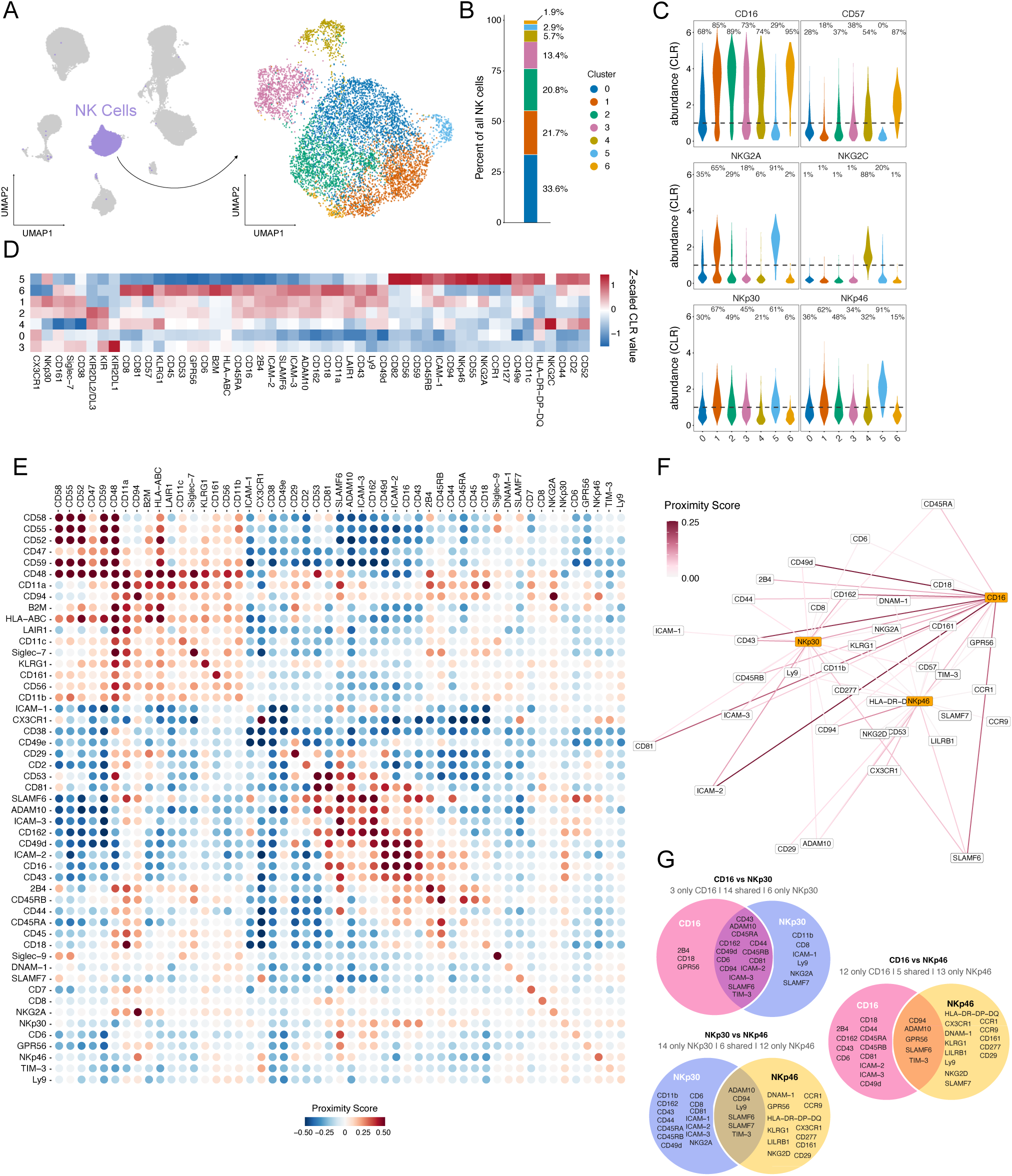
Profiling of the NK cell compartment. (**A**) UMAP projection of re-clustered NK cells identifying distinct subpopulations. (**B**) Relative frequency (%) of identified NK cell subpopulations within the total NK cell compartment. (**C**) Distribution of protein expression levels across NK cell subpopulations; percentages exceeding the detection threshold are indicated. (**D**) Heatmap of protein abundance, expressed as Z-scaled centered log-ratio (CLR) values, for the most variable proteins across identified subpopulations. (**E**) Median per-patient mean proximity scores (log2 ratio) for relevant proteins within NK cell cluster 1. (**F**) Colocalization network of cluster 1, subsetted to the activating receptors CD16, NKp30, and NKp46 and their positively colocalizing partners. (**G**) Pairwise overlap of proteins colocalizing with CD16, NKp30, and NKp46 receptors.

Next, detailed analysis of the membrane organization within the NK cell compartment was performed, focusing on the three largest NK cell clusters (clusters 0-2) (Figure 5E, Figure S5J-K). All clusters demonstrated highly ordered membrane structures containing distinct complexes and multiprotein domains. For instance, cluster 1 was characterized by co-expression of both natural cytotoxicity receptors and Antibody Dependent Cellular Cytotoxicity (ADCC)-inducing CD16, representing an interesting subset for receptor characterization, as CD16 and natural cytotoxicity receptors have been shown to have different functional thresholds and their co-engagement can increase functional responses^23,24^. Within this cluster, a focused analysis on the organization of CD16, NKp30 and NKp46 revealed both shared and receptor-specific interaction partners of the receptors (Figure 5F-G). Notably, CD16 and NKp30 displayed a highly similar interaction landscape, whereas NKp46 mostly presented a more unique profile (Figure 5G). This suggests that the natural cytotoxicity receptors NKp30 and NKp46 are differentially regulated within the plasma membrane.

Together, although PNA resolved considerable heterogeneity in receptor repertoire and receptor organization within circulating NK cells, these features were largely independent of clinical outcome.

### Membrane organization of CD4 T cells provides a dominant predictive signal of response to anti-PD-1 therapy

Altogether, these findings demonstrate significant response-associated differences in immune cell composition, protein abundance, and membrane organization. However, these analyses generally considered each feature independently and may therefore overlook coordinated patterns distributed across multiple markers. To determine whether complex immune profiles could predict clinical outcome, Stabl Lasso was applied, as a feature selection framework designed to identify robust multivariable signatures while minimizing overfitting^25^ (Figure 6A-B). Models were trained to classify patients as responders (OR) or non-responders (non-OR) using either protein abundance or protein proximity measurements from individual immune cell compartments.

**Figure 6.**
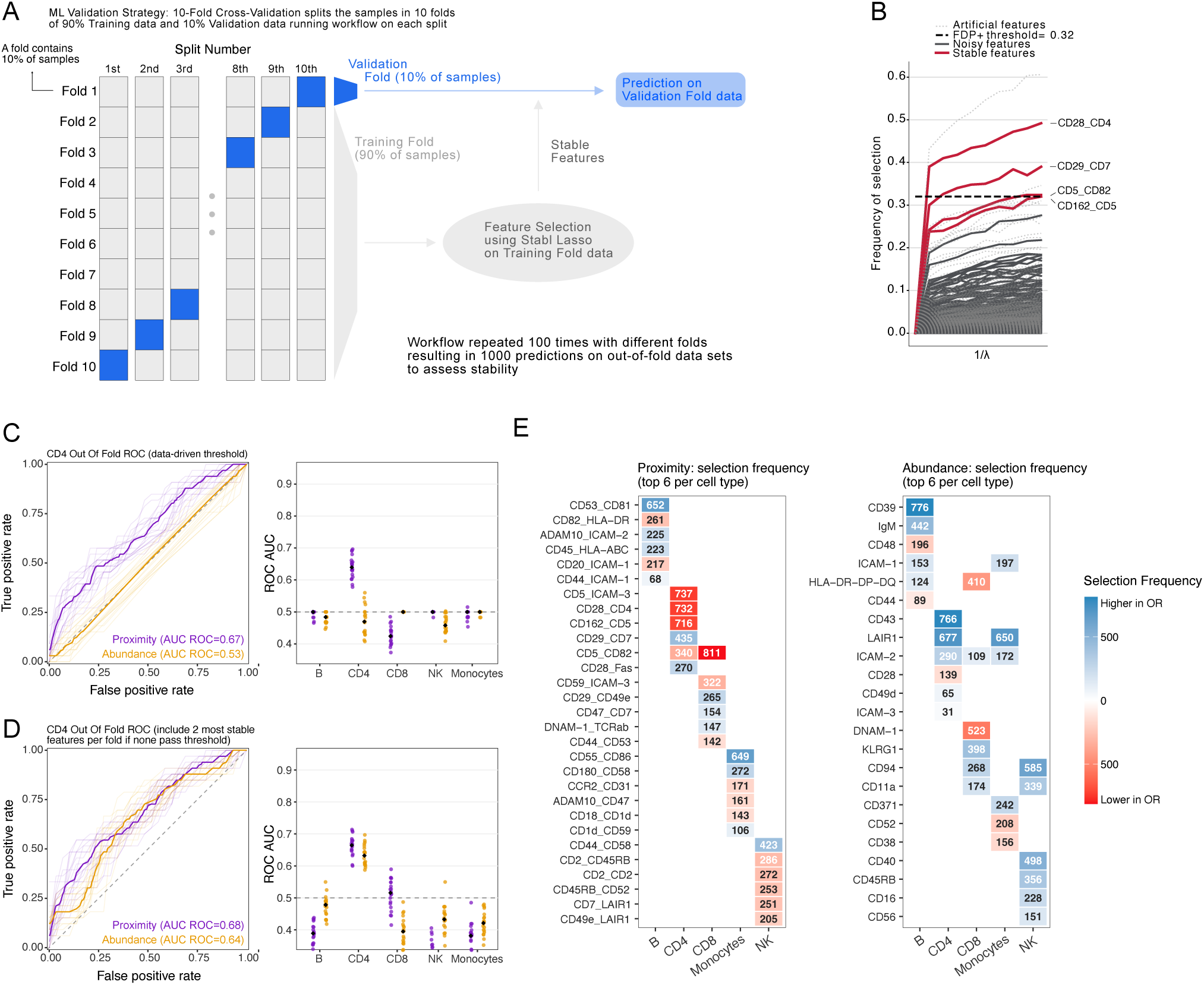
Stabl Lasso feature selection and model performance. (**A**) Overview of the Stabl Lasso validation workflow. (**B**) Example of a single prediction fold stability path in CD4 T cells plotting the mean stability of features across subsamples as the regularization parameter is varied; stability is defined as the proportion of times a feature is selected when trained on subsets of the data and highlighted features indicate stable selection above threshold θ. (**C**) Out-of-fold receiver operating characteristic (ROC) curves for CD4 T cell models, applying a rigorous artificial feature-derived threshold. Shown in bold is the median ROC curve across 20 seeds (left). Distribution of ROC area under the curve (AUC) values across 20 independent runs for major cell types based on abundance or proximity data(right). (**D**) Out-of-fold ROC curves for CD4 T cell models using a relaxed feature selection threshold (top 2 features selected). Distribution of ROC AUC values for 20 runs using a relaxed threshold (right). (**E**) Feature selection frequency across 1,000 model iterations; values represent selection counts using the artificial feature threshold for proximity (left) and abundance (right) data. The 6 most frequent features per cell type are shown.

Although predictive performance varied substantially across cell types (Figure 6C-D), among all compartments examined, CD4 T cells yielded the strongest predictive signal, whereas B cells, CD8 T cells, monocytes, and NK cells showed limited predictive power, further emphasizing the dominant contribution of the CD4 compartment to treatment outcome in this cohort.

An in-depth comparison of abundance- and proximity-based models further revealed that protein organization provided greater predictive power than protein abundance. Using the standard data-driven Stabl Lasso selection threshold, proximity-based CD4 models achieved the highest predictive performance, with a median ROC AUC of 0.67 compared with 0.53 for abundance-based models (Figure 6C-D). Applying a more relaxed selection strategy improved the performance of abundance-based models (ROC AUC = 0.64), although proximity features remained the strongest predictors of clinical outcome (ROC AUC = 0.68).

To identify the features contributing most consistently to these predictions, the frequency with which individual variables were selected across 1,000 model iterations with different training data sets was examined (Figure 6E). Once again, the resulting signatures were dominated by CD4-derived features, particularly proximity measurements, and included interactions between CD5, CD28, CD162, ICAM-3, CD4, and CD82. Notably, several of the features were not individually significant in the differential analyses, indicating that a broader restructuring of CD4 T cell membrane consisting of modest but coordinated changes correlates with treatment outcome. These findings highlight the value of multivariate modeling for identifying response-associated immune states that cannot be captured by single-feature analyses alone.

Taken together, these results establish the CD4 T cell compartment as the dominant source of response-associated information in this cohort. While both abundance and proximity measurements contributed to outcome prediction, membrane organization consistently provided the strongest predictive signal, suggesting that clinically relevant immune states are captured by the spatial organization of cell surface proteins.

## Discussion

Despite the transformative impact of immune checkpoint inhibitors, only a subset of patients achieves durable clinical responses, highlighting the need for reliable biomarkers that can guide treatment selection and spare non-responding patients from ineffective therapy and unnecessary toxicity^3^. Although extensive efforts have been devoted to identifying predictive immune biomarkers, these have been almost exclusively focused on immune cell frequencies, transcriptional profiles, or protein abundance^26^. In contrast, the spatial organization of proteins at the cell surface has received little attention despite its fundamental role in regulating immune cell communication and signaling. To our knowledge, this is the first study to investigate whether cell surface protein organization is associated with clinical response to anti-PD-1 therapy. By applying the Proximity Network Assay (PNA) to pretreatment PBMCs from patients with advanced NSCLC, we demonstrated that response-associated immune states are encoded not only by differences in protein abundance but also by changes in cell surface architecture. These alterations were most pronounced within the CD4 T cell compartment, where both protein expression and their spatial organization were independently associated with treatment outcome.

It is notable that, in this study, the strongest response-associated alterations were observed within CD4 T cells, rather than cytolytic lymphocytes such as CD8 T cells and NK cells. In fact, as CD8 T cells are widely regarded as the primary effector population responsible for tumor cell killing following anti-PD-1 therapy, many studies have therefore focused on their abundance, activation state, or clonal expansion^27–29^ to predict response. On the contrary, we found that circulating CD8 T cells, despite being relatively fewer in responding patients, possibly because of increased infiltration into the tumor mass as previously reported^30^, exhibited remarkably limited phenotypic variation between response groups. On the other hand, the CD4 compartment displayed both compositional and molecular alterations associated with clinical outcome. Within this compartment, responders showed elevated expression of several adhesion molecules, suggesting a heightened migratory immune state prior to treatment, possibly enabling increased potential for tumor infiltration. Additionally, in accordance with previous reports, monocyte populations displayed correlation to clinical response^31^, with non-classical monocytes being more frequent in responding patients. Within this compartment, complete responders (CR) additionally displayed reduced platelet-monocyte aggregates, suggesting an altered activation of platelet-coated monocytes possibly leading to impaired inflammatory responses.

Beyond conventional phenotypic profiling, our data revealed that membrane organization represents a key additional layer of immune cell regulation associated with response to immune checkpoint therapy. Differential proximity analysis identified altered organization of a CD5-containing protein domain at the surface of CD4 T cells. Notably, many of the proteins in this domain, including CD162, CD11a, CD43, CD44, CD28, and SLAMF6, are known to participate in cell communication, migration, immune synapse formation, or regulation of T cell activation^32–34^. Increased colocalization within this module was associated with reduced response and overall survival, indicating potential clinical relevance. Importantly, increased cis-interactions of one of these domain proteins, SLAMF6, was recently associated with loss of T cell function in cancer^35^. Rather than observing isolated changes in individual protein pairs, we detected coordinated remodeling of an interconnected membrane domain, suggesting that response-associated immune states may be reflected in higher-order membrane architecture.

The biological significance of this organizational signature remains to be determined. For instance, CD5 functions as an important regulator of T cell receptor signaling and has been implicated in tuning activation thresholds and maintaining immune homeostasis^36,37^. The observation that CD5 exhibited altered spatial relationships with multiple classes of proteins raises the possibility that membrane organization influences how T cells integrate signals from antigen receptors, co-stimulatory receptors, and adhesion pathways. Such mechanisms could affect T cell activation, trafficking, or responsiveness to checkpoint blockade. While our study does not establish causality, it highlights membrane organization as a potentially important and previously underappreciated feature of immune cell biology in cancer immunotherapy.

The observation that protein organization outperformed protein abundance in predictive modeling further supports this concept. Traditional immune profiling approaches primarily quantify single altered features, implicitly assuming that proteins function independently. However, cell surface proteins operate within highly organized molecular networks whose spatial arrangement can profoundly influence signaling and cellular behavior^38,39^. Consistent with this, proximity-based models provided greater predictive performance than abundance-based models and identified response-associated signatures that extended beyond individually significant markers. Although these findings should be considered hypothesis generating given the relatively modest cohort size, they demonstrate the value of incorporating cell surface protein organization into immune profiling. Validation in larger, independent patient cohorts will be essential to confirm the robustness and clinical utility of the identified signatures.

In conclusion, responders showed compositional and phenotypical variation associated with outcome in CD4 T cells and more specifically in a newly described CD5-containing architectural domain. To our knowledge this specific architecture has not been previously described and warrants further investigation. Altogether, these findings also suggest that biomarker discovery efforts should consider how proteins are spatially organized at the cell surface. Incorporating this additional layer of biological information has the potential to uncover entirely novel biomarkers and improve our understanding of the immune mechanisms underlying successful cancer immunotherapy.

## Methods

### Sample Collection

PBMC were isolated from the peripheral blood of 62 NSCLC patients by density gradient centrifugation, layering collected blood on top of Ficoll-Paque (Cytiva AB) at 2:1 ratio and centrifuging for 20 minutes at 2000rpm. Then, following one wash in PBS, collected PBMC were added to 5mL of a red blood cell-lysis solution (NH_4_Cl and NaHCO_3_ in ddH_2_O + 2% EDTA 0.5M) and incubated for 5 minutes at 37 degrees. Finally, cells were resuspended at 2-10 million/vial in heat-inactivated FBS supplemented with 10% DMSO and stored long term in liquid nitrogen.

### Patients, treatments and endpoints

PMBCs were obtained from a cohort of 62 patients with advanced NSCLC who received anti-PD1-therapy at Karolinska University Hospital between 2021 and 2025 (Table 1). Patients receiving high dose corticosteroids were excluded. Anti-PD1-therapy was administered as monotherapy or in combination with an initial four cycles of chemotherapy. The anti-PD1-therapy continued until disease progression, intolerable adverse events or up to 2 years. Patient and tumor characteristics were gathered from the electronic medical records. Clinical endpoints consisted of tumor response and survival outcomes. Objective response (complete or partial response, stable disease or progressive disease) was assessed by retrospectively reviewing the radiological images and reports according to RECIST 1.1 criteria. Progression-free survival (PFS) and overall survival (OS) were calculated from the time of initiation of treatment to the occurrence of disease progression or death, respectively. Patients not experiencing disease progression or death during follow-up were censored at the last contact date. No cases were recorded as lost to follow up.

### Proximity Network Assay

Samples were processed in two batches, with OR and non-OR samples represented in each batch. In short, PBMCs were thawed, washed and fixed using 1% PFA, 15 minutes at room temperature. Fixation was quenched using 135 mM glycine and washed. The cells were either processed in PNA immediately (16 samples) or frozen and stored using 95% FBS + 5% DMSO, and thawed and processed a few weeks post freezing (46 samples). PNA was carried out according to the Pixelgen Proxiome Kit user manual. In short, cells from each sample were labeled with a cocktail containing 158 oligoconjugated antibodies. Proximity networks were generated and subsequent sequences were amplified using PCR, purified and quantified. The first batch of samples was sequenced in a 10B flow cell using the Illumina NovaSeq X Plus sequencing system (PE-100). The second batch of samples was sequenced using a DNBSEQ-T7 platform (MGI Tech, Shenzhen, China) according to a paired-end sequencing strategy (Read 1: 45bp, Read 2: 79bp).

### Data Preprocessing

Data processing for the PNA sequencing data was performed using the open-source pixelator pipeline (nf-core/pixelator, v2.3.1). Briefly, raw reads were subjected to quality control and filtering, followed by amplicon assembly. These amplicons represent a link between two proteins and were then grouped by marker assignment and error-corrected to construct sample-wide edge lists and graphs. Individual cells were identified using Leiden community detection, with only graphs exceeding a node size threshold of 8,000 retained. Spatial information is calculated for each individual cellular component and compiled, along with additional data, in a single sample-specific PXL file. These PXL files are then used as input for downstream analysis in R using pixelatorR v0.18.2. https://github.com/PixelgenTechnologies/pixelatorR).

### Downstream Analysis

#### Proximity Score

To assess the degree of spatial association between protein markers, a metric termed the Proximity Score, which is based on join count statistics, was utilized. This metric compares the observed number of direct edges between two proteins within a component graph (the join count *x*) to the expected number of edges based on the respective protein abundances (*E*). The Proximity Score (*PS*) is then expressed as the *log*2 ratio:

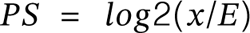

Calculating the Proximity Score in this manner effectively provides an abundance-normalized measure, allowing for the comparison of spatial associations across varying ranges of protein expression levels.

#### Annotation

Single-cell annotation was conducted through an integrative approach combining label transfer and gating-based strategies. Initially, cell protein abundance profiles were mapped to a previously annotated reference PBMC PNA dataset to assign cells to major lineages. Subsequently, marker-specific gating was employed to refine subpopulation annotations.

#### Re-clustering strategy

Each major immune lineage was subset and re-clustered independently: the top 30 principal components were computed based on a curated lineage-specific marker panel, integrated across samples with Harmony, and embedded with UMAP from the Harmony components. PCA and neighborhood parameters were tuned per lineage to account for differences in cell number and panel size. For the NK cells, re-clustering was performed at res 0.25 using the Louvain algorithm.

#### Marker pair selection

For visualizations involving Proximity Scores - including colocalization heatmaps and network representations - proteins were filtered based on minimum abundance and frequency thresholds within the relevant cell population. Cut-offs were determined using isotype control data and background signal levels observed in negative populations.

#### Differential Testing

Differential analyses were performed on sample-level pseudo-bulked proximity scores within each cell type (mean log2 ratio for proximity and mean CLR values for abundance), testing associations with clinical outcome using a linear model and batch included as a covariate. P-values were adjusted for multiple testing across features within each cell type using the Benjamini–Hochberg false discovery rate procedure. Effect sizes represent batch-adjusted differences in means between outcomes. Differential cell-type frequency analysis was performed with the propeller function from the R package speckle (v0.99.7), which computes per-sample cell-type proportions, applies a logit transformation, and tests for differences in composition between clinical response groups using empirical Bayes–moderated linear models with a mean–variance trend.

#### Survival analysis

Within each batch, patients were ranked by mean abundance or proximity score and split at the median; survival was then compared between the lower and upper halves. Kaplan–Meier curves were estimated with *survfit(Surv(os_days, os_censor) ∼ group)* using the survival (v3.8.6) and survminer (v0.5.2) R packages. Group differences were tested with the log-rank test. A p value <0.05 was considered statistically significant. In addition, a continuous Cox proportional hazards model was fitted using all samples. Pseudo-bulked Proximity Scores were z-scored within the batch and modeled as *Surv(os_days, os_censor) ∼ proximity_z*. The hazard ratio (HR) per 1 standard deviation increase in proximity, with 95% confidence interval and p-value, was displayed in a forest plot on a log scale (reference HR = 1).

#### Monocyte-platelet aggregation

To segment cell graphs into host-cell and platelet compartments, protein signatures distinguishing the host compartment from platelets were defined. Non-negative matrix factorization (NMF) was applied to the protein abundance profiles of the two populations (host and platelet) to learn two compartment-specific weight profiles. For each single-cell graph, nodes were assigned to a compartment based on their local protein composition. Because individual nodes carry only a single protein identity, each node’s profile was first aggregated over its k-hop graph neighborhood, pooling counts from directly and indirectly connected nodes so that the local molecular environment, rather than a single noisy measurement, drives classification. Neighborhood-aggregated protein profiles were projected onto the two compartment signatures using a non-negative projection, yielding per-node compartment scores. Nodes were then assigned to a compartment by thresholding the normalized score with a data-driven cutoff derived from a two-center k-means clustering of the score distribution. Node assignments were subsequently spatially smoothed. Boundary nodes between the two compartments were labeled as interface nodes.

#### Multivariable analysis

For each immune cell population, two modalities were compared as predictors of clinical response (OR vs non-OR): proximity (sample-level pseudo-bulked mean Proximity Scores for marker pairs within the cell type) and abundance (sample-level pseudo-bulked mean CLR values within the cell type). All predictors were z-scored within the batch prior to modeling. Feature selection and prediction were performed using Stabl Lasso. In Stabl, the data are repeatedly subsampled and fitted with logistic Lasso across a range of λ penalties. Artificial null features unrelated to the outcome are injected to estimate a surrogate false discovery proportion (FDP⁺), and a data-driven reliability threshold (θ) is selected as the frequency cutoff that minimizes FDP⁺; features with selection frequency above θ constitute the final sparse and stable marker set. Model performance was assessed using cross-validated out-of-fold predictions using standard logistic regression with these stable features, from which ROC curves and AUC were computed. Crucially, prediction is thus always performed on samples that were not used during the training step. Analyses were repeated across 20 random seeds (each 5 × 10-fold cross-validation repeats, yielding 1,000 resampling runs for stability summaries). To identify the features contributing most consistently to prediction, the frequency with which individual features were selected across the 1,000 model runs was examined.

## Declarations

This study was approved by the Swedish Ethical Review Authority (#2019-00093 and #2025-01215-02). VvH, HvO and SF are full-time employees of Pixelgen Technologies. All other authors declare no competing interests.

## Acknowledgements

This study was supported by the Cancer Research Foundations of Radiumhemmet (#201103, #231123, and #241293), The Swedish Cancer Society (#CAN24 3655 Pj and #CAN23 2808 Fk), The Swedish Research Council (#2022-00723), Karolinska Institutet (#2023-01411), The Sjöberg Foundation, and Pixelgen Technologies.

## Author contributions

AL, RC, and SF conceived the project. RC, YingY, VvH, HvO, MG, NB, MC, LBG, and YimingY, carried out experiments. RC, YingY, VvH, HvO, AP, YimingY, LL, and LdP analyzed the data. AP, LL, SF, and LdP provided key reagents, and patient samples and data. AL, YingY, LL, LdP, MJ, KK, and SF obtained funding and supervised the work. RC, VvH, HvO, and AL wrote the manuscript with input from all authors. All authors reviewed and approved the final manuscript.

## References

1. West H, McCleod M, Hussein M, et al. Atezolizumab in combination with carboplatin plus nab-paclitaxel chemotherapy compared with chemotherapy alone as first-line treatment for metastatic non-squamous non-small-cell lung cancer (IMpower130): a multicentre, randomised, open-label, phase 3 trial. Lancet Oncol. 2019;20(7):924–937. doi:10.1016/S1470-2045(19)30167-6

2. Sezer A, Kilickap S, Gümüş M, et al. Cemiplimab monotherapy for first-line treatment of advanced non-small-cell lung cancer with PD-L1 of at least 50%: a multicentre, open-label, global, phase 3, randomised, controlled trial. Lancet. 2021;397(10274):592–604. doi:10.1016/S0140-6736(21)00228-2

3. Mok TSK, Wu YL, Kudaba I, et al. Pembrolizumab versus chemotherapy for previously untreated, PD-L1-expressing, locally advanced or metastatic non-small-cell lung cancer (KEYNOTE-042): a randomised, open-label, controlled, phase 3 trial. Lancet. 2019;393(10183):1819–1830. doi:10.1016/S0140-6736(18)32409-7

4. de Castro G, Kudaba I, Wu YL, et al. Five-Year Outcomes With Pembrolizumab Versus Chemotherapy as First-Line Therapy in Patients With Non–Small-Cell Lung Cancer and Programmed Death Ligand-1 Tumor Proportion Score ≥ 1% in the KEYNOTE-042 Study. J Clin Oncol. 2023;41(11):1986–1991. doi:10.1200/JCO.21.02885

5. Reck M, Rodríguez-Abreu D, Robinson AG, et al. Five-Year Outcomes With Pembrolizumab Versus Chemotherapy for Metastatic Non-Small-Cell Lung Cancer With PD-L1 Tumor Proportion Score ≥ 50. J Clin Oncol Off J Am Soc Clin Oncol. 2021;39(21):2339–2349. doi:10.1200/JCO.21.00174

6. Kilickap S, Baramidze A, Sezer A, et al. Cemiplimab Monotherapy for First-Line Treatment of Patients with Advanced NSCLC With PD-L1 Expression of 50% or Higher: Five-Year Outcomes of EMPOWER-Lung 1. J Thorac Oncol Off Publ Int Assoc Study Lung Cancer. 2025;20(7):941–954. doi:10.1016/j.jtho.2025.03.033

7. Garon EB, Hellmann MD, Rizvi NA, et al. Five-Year Overall Survival for Patients With Advanced Non‒Small-Cell Lung Cancer Treated With Pembrolizumab: Results From the Phase I KEYNOTE-001 Study. J Clin Oncol Off J Am Soc Clin Oncol. 2019;37(28):2518–2527. doi:10.1200/JCO.19.00934

8. Biton J, Mansuet-Lupo A, Pécuchet N, et al. TP53, STK11, and EGFR Mutations Predict Tumor Immune Profile and the Response to Anti-PD-1 in Lung Adenocarcinoma. Clin Cancer Res Off J Am Assoc Cancer Res. 2018;24(22):5710–5723. doi:10.1158/1078-0432.CCR-18-0163

9. Liu Z, Yang Z, Wu J, et al. A single-cell atlas reveals immune heterogeneity in anti-PD-1-treated non-small cell lung cancer. Cell. 2025;188(11):3081–3096.e19. doi:10.1016/j.cell.2025.03.018

10. Augustyn A, Adams DL, He J, et al. Giant Circulating Cancer-Associated Macrophage-Like Cells Are Associated With Disease Recurrence and Survival in Non-Small-Cell Lung Cancer Treated With Chemoradiation and Atezolizumab. Clin Lung Cancer. 2021;22(3):e451–e465. doi:10.1016/j.cllc.2020.06.016

11. Inomata M, Matsumoto M, Takata N, et al. Peripheral CD4 memory T cells predict the efficacy of immune checkpoint inhibitor therapy in patients with non-small cell lung cancer. Sci Rep. 2023;13(1):10807. doi:10.1038/s41598-023-37736-3

12. Cao J, Zhang Y, Guo S, et al. Immune biomarkers in circulating cells of NSCLC patients can effectively evaluate the efficacy of chemotherapy combined with anti-PD-1 therapy. Front Immunol. 2025;16:1521708. doi:10.3389/fimmu.2025.1521708

13. Kim GD, Shin SI, Sun P, et al. Single-cell RNA sequencing of baseline PBMCs predicts ICI efficacy and irAE severity in patients with NSCLC. J Immunother Cancer. 2025;13(5). doi:10.1136/jitc-2025-011636

14. van Deventer S, Arp AB, van Spriel AB. Dynamic Plasma Membrane Organization: A Complex Symphony. Trends Cell Biol. 2021;31(2):119–129. doi:10.1016/j.tcb.2020.11.004

15. Lacouture C, Chaves B, Guipouy D, et al. LFA-1 nanoclusters integrate TCR stimulation strength to tune T-cell cytotoxic activity. Nat Commun. 2024;15(1):407. doi:10.1038/s41467-024-44688-3

16. Salavessa L, Lagache T, Malardé V, et al. Cytokine receptor cluster size impacts its endocytosis and signaling. Proc. Natl. Acad. Sci. U.S.A. 2021;118 (37) e2024893118, 10.1073/pnas.2024893118

17. Sarén T, Saronio G, Marti Torrell P, et al. Complementarity-determining region clustering may cause CAR-T cell dysfunction. Nat Commun. 2023;14(1):4732. doi:10.1038/s41467-023-40303-z

18. Philips EA, Liu J, Kvalvaag A, et al. Transmembrane domain-driven PD-1 dimers mediate T cell inhibition. Sci Immunol. 2024;9(93):eade6256. doi:10.1126/sciimmunol.ade6256

19. Li B, Lu Y, Zhong MC, et al. Cis interactions between CD2 and its ligands on T cells are required for T cell activation. Sci Immunol. 2022;7(74):eabn6373. doi:10.1126/sciimmunol.abn6373

20. Zhao Y, Caron C, Chan YY, et al. Cis-B7:CD28 Interactions at Invaginated Synaptic Membranes Activate CD28 and Promote CD8+ T Cell Function and Anti-Tumor Immunity. Immunity. 2023;56(6):1187–1203.e12. doi:10.1016/j.immuni.2023.04.005

21. Single-Cell Protein Interactomes by the Proximity Network Assay | bioRxiv. Accessed August 18, 2026. https://www.biorxiv.org/content/10.1101/2025.06.19.660329v1

22. Rebuffet L, Melsen JE, Escalière B, et al. High-dimensional single-cell analysis of human natural killer cell heterogeneity. Nat Immunol. 2024 Aug;25(8):1474–1488. doi: 10.1038/s41590-024-01883-0

23. Bryceson YT, March ME, Ljunggren HG, et al. Synergy among receptors on resting NK cells for the activation of natural cytotoxicity and cytokine secretion. Blood. 2006 Jan 1;107(1):159–66. doi: 10.1182/blood-2005-04-1351.

24. Gauthier L, Morel A, Anceriz N, et al. Multifunctional Natural Killer Cell Engagers Targeting NKp46 Trigger Protective Tumor Immunity. Cell. 2019;177(7):1701–1713.e16. doi:10.1016/j.cell.2019.04.041

25. Hédou J, Marić I, Bellan G, et al. Discovery of sparse, reliable omic biomarkers with Stabl. Nat Biotechnol. 2024;42(10):1581–1593. doi:10.1038/s41587-023-02033-x

26. Tostes K, Siqueira AP, Reis RM, Leal LF, Arantes LMRB. Biomarkers for Immune Checkpoint Inhibitor Response in NSCLC: Current Developments and Applicability. Int J Mol Sci. 2023;24(15):11887. doi:10.3390/ĳms241511887

27. Kamphorst AO, Pillai RN, Yang S, et al. Proliferation of PD-1+ CD8 T cells in peripheral blood after PD-1–targeted therapy in lung cancer patients. Proc Natl Acad Sci U S A. 2017;114(19):4993–4998. doi:10.1073/pnas.1705327114

28. Thommen DS, Koelzer VH, Herzig P, et al. A transcriptionally and functionally distinct PD-1+ CD8+ T cell pool with predictive potential in non-small cell lung cancer treated with PD-1 blockade. Nat Med. 2018;24(7):994–1004. doi:10.1038/s41591-018-0057-z

29. Kim H, Park S, Han KY, et al. Clonal expansion of resident memory T cells in peripheral blood of patients with non-small cell lung cancer during immune checkpoint inhibitor treatment. J Immunother Cancer. 2023;11(2):e005509. doi:10.1136/jitc-2022-005509

30. Niemeĳer ALN, Sahba S, Smit EF, Lissenberg-Witte BI, de Langen AJ, Thunnissen E. Association of tumour and stroma PD-1, PD-L1, CD3, CD4 and CD8 expression with DCB and OS to nivolumab treatment in NSCLC patients pre-treated with chemotherapy. Br J Cancer. 2020;123(3):392–402. doi:10.1038/s41416-020-0888-5

31. Ohkuma R, Fujimoto Y, Ieguchi K, et al. Monocyte subsets associated with the efficacy of anti-PD-1 antibody monotherapy. Oncol Lett. 2023;26(3):381. doi:10.3892/ol.2023.13967

32. Walling BL, Kim M. LFA-1 in T Cell Migration and Differentiation. Front Immunol. 2018;9:952. doi:10.3389/fimmu.2018.00952

33. Ali AJ, Abuelela AF, Merzaban JS. An Analysis of Trafficking Receptors Shows that CD44 and P-Selectin Glycoprotein Ligand-1 Collectively Control the Migration of Activated Human T-Cells. Front Immunol. 2017;8. doi:10.3389/fimmu.2017.00492

34. Ciesielska-Figlon K, Lisowska KA. The Role of the CD28 Family Receptors in T-Cell Immunomodulation. Int J Mol Sci. 2024;25(2):1274. doi:10.3390/ĳms25021274

35. Li B, Zhong MC, Galindo CC, et al. SLAMF6 as a drug-targetable suppressor of T cell immunity against cancer. Nature. 2026;652(8110):722–730. doi:10.1038/s41586-026-10106-5

36. Tarakhovsky A, Kanner SB, Hombach J, et al. A role for CD5 in TCR-mediated signal transduction and thymocyte selection. Science. 1995;269(5223):535–537. doi:10.1126/science.7542801

37. Peña-Rossi C, Zuckerman LA, Strong J, et al. Negative regulation of CD4 lineage development and responses by CD5. J Immunol. 1999;163(12):6494–6501.

38. Chao Z, Mei Q, Yang C, et al. Immunological synapse: structures, molecular mechanisms and therapeutic implications in disease. Signal Transduct Target Ther. 2025;10(1):254. doi:10.1038/s41392-025-02332-6

39. Sezgin E, Levental I, Mayor S, Eggeling C. The mystery of membrane organization: composition, regulation and physiological relevance of lipid rafts. Nat Rev Mol Cell Biol. 2017;18(6):361–374. doi:10.1038/nrm.2017.16

