## Supplementary data for "CD4 T cell surface architecture is associated with response to anti-PD-1 therapy in non-small cell lung cancer"

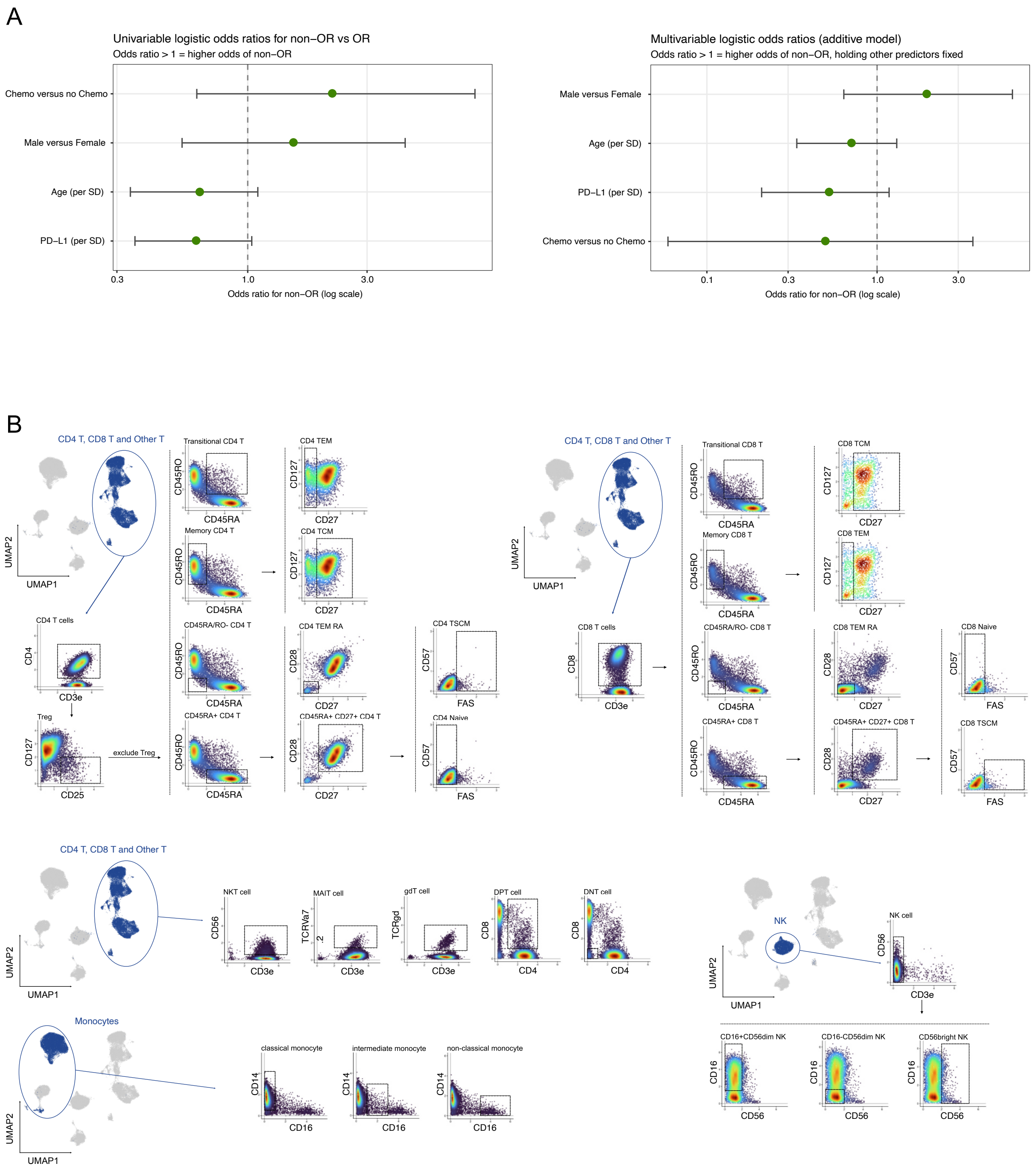


***Figure 1 Supplement.*** *(**A**) Logistic regression of clinical predictors to best response. Values above the dotted line indicate higher odds of non-response; each predictor fit separately (left) and predictors fit together in one additive model (right). (**B**) Gating strategy for the major cell lineages.*

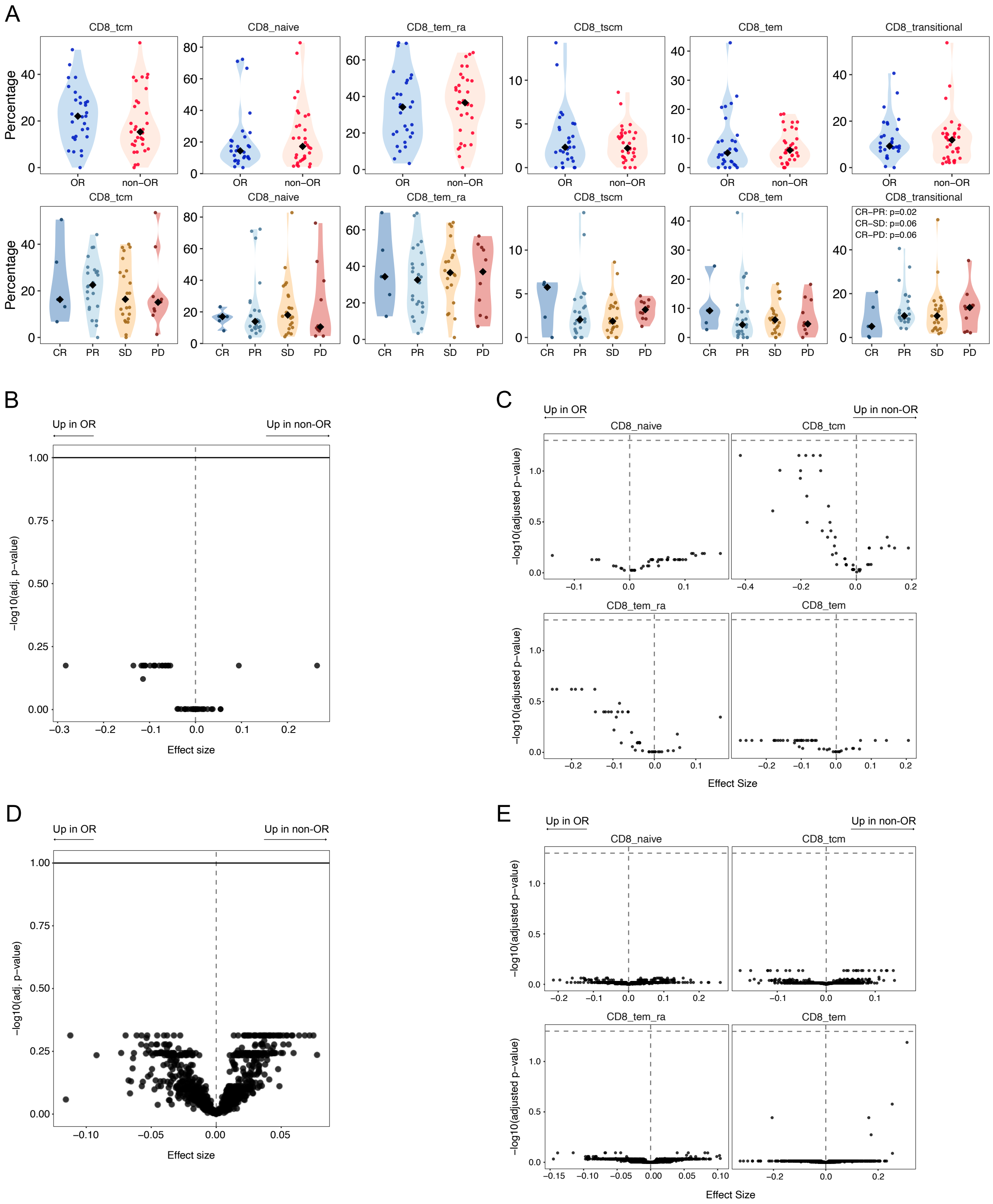


***Figure 2 Supplement. Differential analyses of the CD8 compartment.*** *(**A**) Quantification of CD8 T cell subpopulation proportions within the total CD8 compartment split by outcome (top row) and clinical response (bottom row). Each dot represents a single patient and p-values < 0.1 are indicated. Volcano plots illustrating differential abundance between OR and non-OR for the whole CD8 T cell population (**B**) or within individual CD8 T cell subsets (**C**). Horizontal line denotes adjusted p-value < 0.05. Volcano plots illustrating differential proximity scores between OR and non-OR for the whole CD8 T cell population (**D**) or within individual CD8 T cell populations (**E**). Horizontal line denotes adjusted p-value < 0.05.*

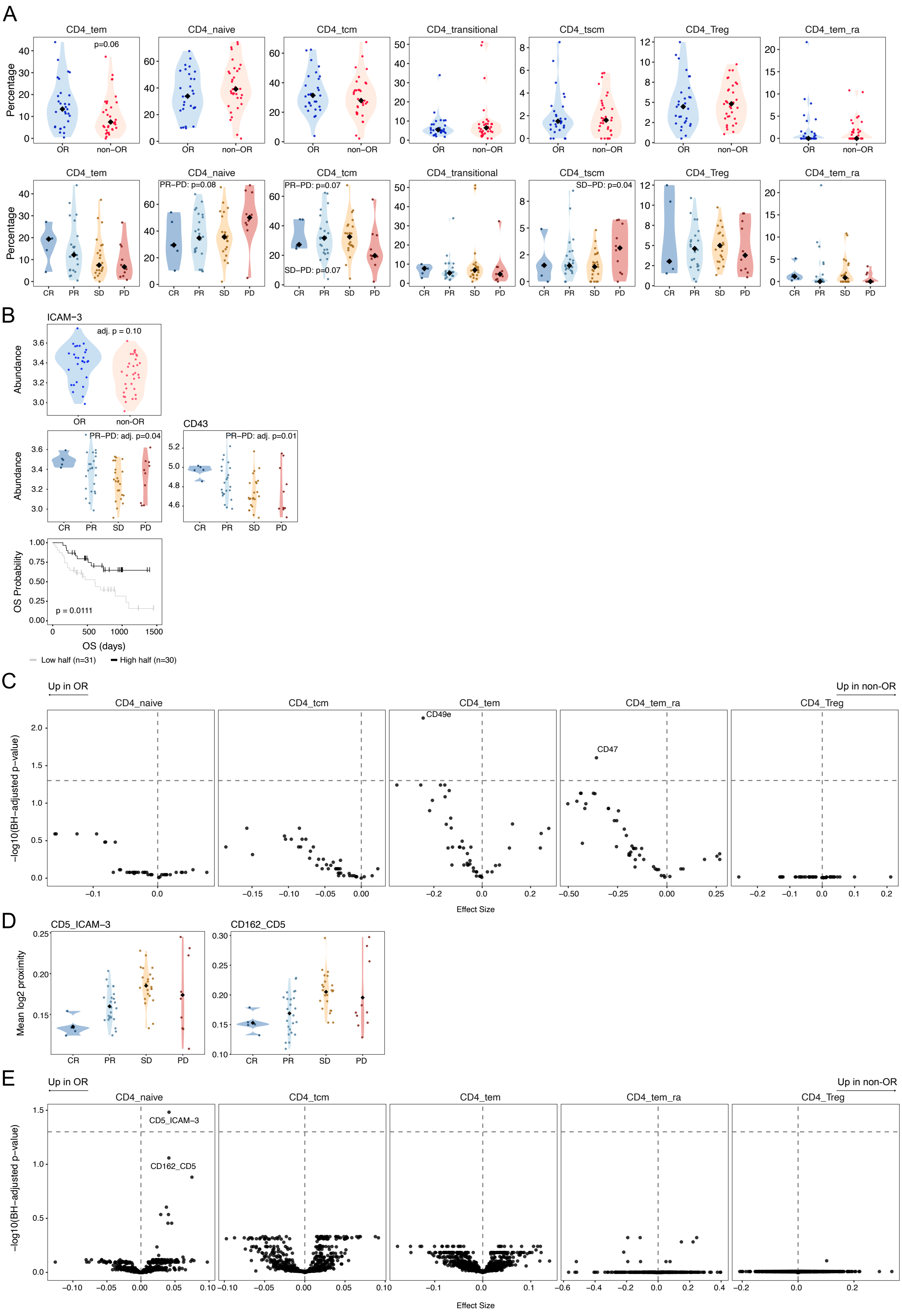

A
CD4_tem
CD4_naive
CD4_tcm
CD4_transitional
CD4_tscm
CD4_Treg
CD4_tem_ra
p=0.06
Percentage
OR
non-OR
PR–PD: p=0.08
PR–PD: p=0.07
SD–PD: p=0.07
SD–PD: p=0.04
CR
PR
SD
PD
B
ICAM−3
adj. p = 0.10
Abundance
CD43
PR–PD: adj. p=0.04
PR–PD: adj. p=0.01
OS Probability
p = 0.0111
OS (days)
Low half (n=31)
High half (n=30)
C
Up in OR
Up in non-OR
CD49e
CD47
−log10(BH-adjusted p-value)
Effect Size
D
CD5_ICAM-3
CD162_CD5
Mean log2 proximity
E
CD5_ICAM-3
CD162_CD5

***Figure 3 Supplement. CD4 T cell subpopulation composition.*** *(**A**) Relative frequency of CD4 T cell subpopulations within the total CD4 compartment split by outcome (top row) and clinical response (bottom row). Each dot represents a single patient and p-values < 0.1 are indicated. (**B**) Abundance distribution of selected markers with each dot representing the mean CLR of a patient; stratified by outcome (top) and clinical category (middle). Kaplan–Meier survival analysis comparing patients stratified by the highest versus lowest half of marker abundance (bottom). (**C**) Volcano plots illustrating differential abundance between OR and non-OR for individual CD4 T cell subsets. Horizontal line denotes adjusted p-value < 0.05. (**D**) Proximity Score distribution of significant markers stratified by clinical category; each dot represents the mean log2 ratio of a patient. (**E**) Volcano plots illustrating differential proximity scores between OR and non-OR within the CD4 T cell subpopulations. Horizontal line denotes adjusted p-value < 0.05.*

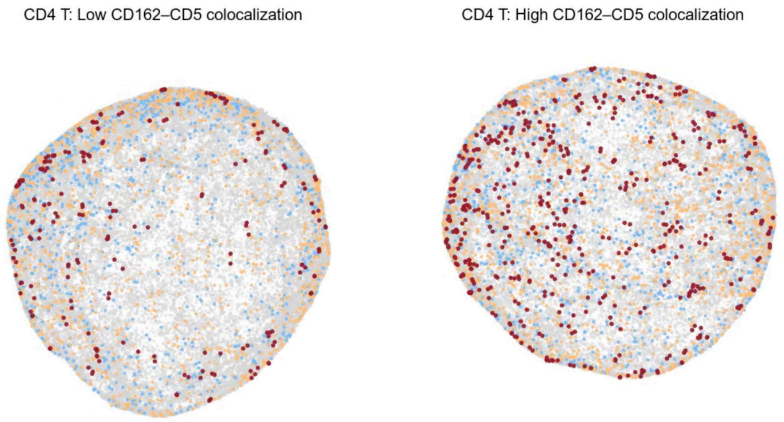


***Supplementary Video 1**. Three-dimensional representation of OR and non-OR CD4 T cells matched for abundance of CD162 (blue) and CD5 (orange) markers. Colocalizing CD162 and CD5 (connected directly by an edge) are represented in red.*

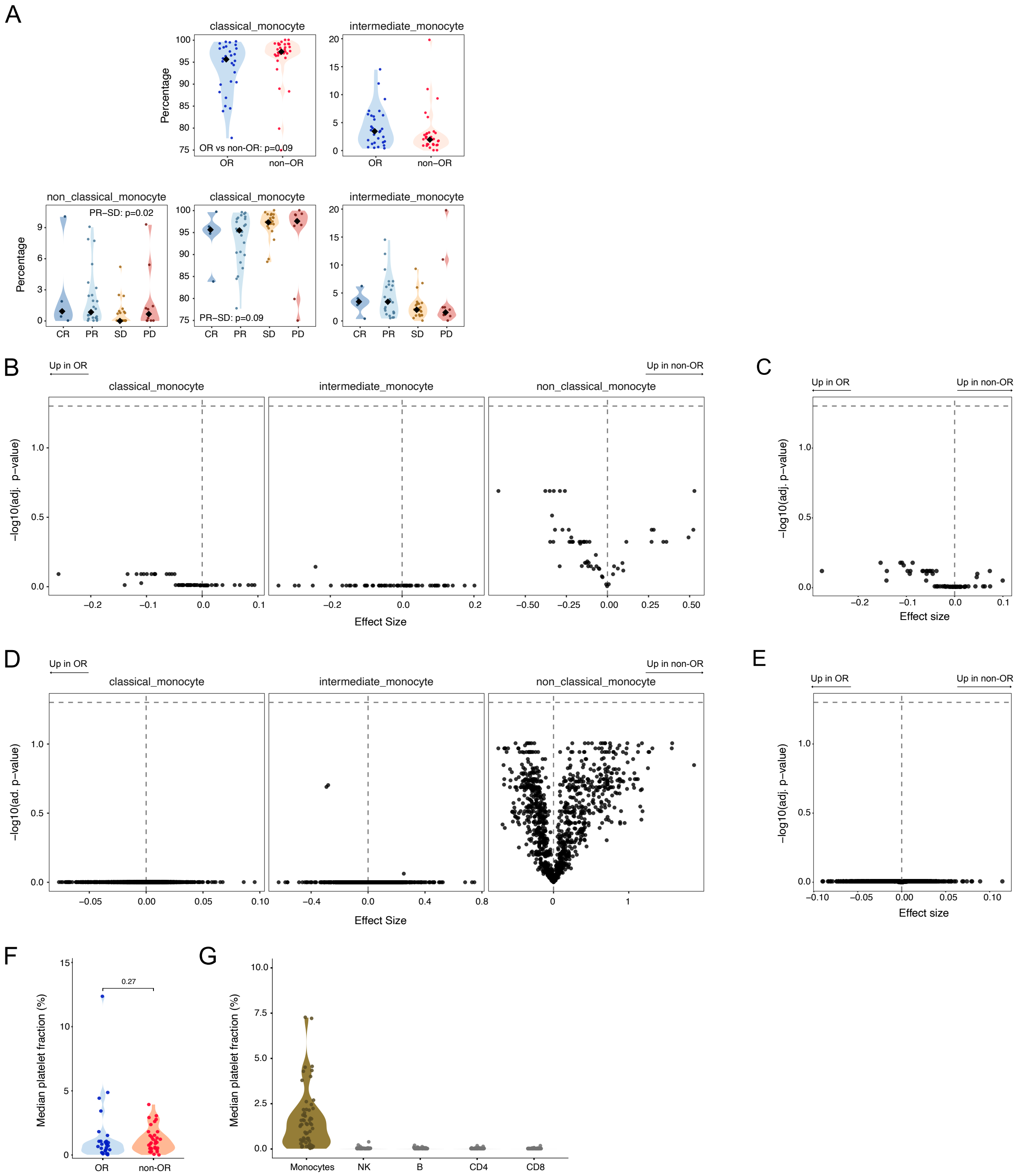


***Figure 4 Supplement. Differential analysis of the monocyte compartment.*** *(**A**) Relative frequency of monocyte subpopulations within the total monocyte compartment; split by outcome (top row) and clinical response (bottom row) and p-values < 0.1 are indicated. Volcano plot illustrating differential abundance between OR and non-OR within the monocyte subpopulations (**B**) and for the whole monocyte compartment (**C**). The horizontal line represents an adjusted p-value of 0.05. Volcano plots illustrating differential proximity between OR and non-OR within the monocyte subpopulations (**D**) and for the whole monocyte compartment (**E**). Horizontal line denotes adjusted p-value < 0.05. (**F**) Quantification of the fraction of nodes assigned to platelets per outcome category. Individual dots indicate the median fraction per patient (based on a subsample of 1000 random monocytes). (**G**) Quantification of the fraction of platelet nodes in a random sample of 500 cells per cell type, each dot represents the median per patient.*

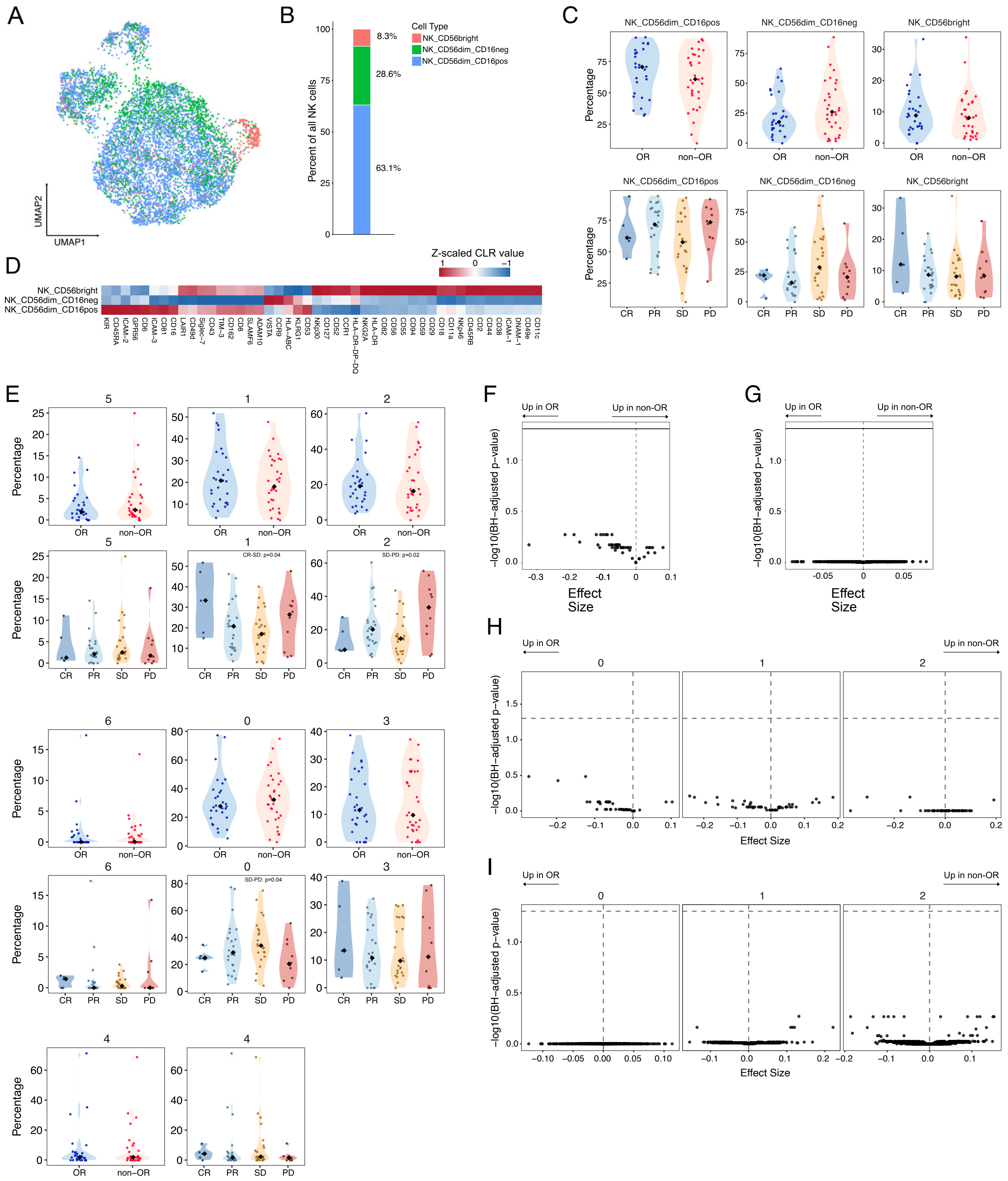
A
UMAP2
UMAP1
B
Percent of all NK cells
Cell Type
NK_CD56bright
NK_CD56dim_CD16neg
NK_CD56dim_CD16pos
8.3%
28.6%
63.1%
C
NK_CD56dim_CD16pos
NK_CD56dim_CD16neg
NK_CD56bright
Percentage
OR
non−OR
CR
PR
SD
PD
D
Z-scaled CLR value
NK_CD56bright
NK_CD56dim_CD16neg
NK_CD56dim_CD16pos
KIR
CD45RA
ICAM-2
GPR56
CD6
ICAM-3
CD81
CD16
LAIR1
CD49d
Siglec-7
CD43
TIM-3
CD162
CD8
SLAMF6
ADAM10
VISTA
CCR9
HLA-ABC
KLRG1
CD53
NKp30
CD127
CD52
CCR1
HLA-DR-DP-DQ
NKG2A
HLA-DR
CD82
CD56
CD55
CD94
CD59
CD29
CD18
CD11a
NKp46
CD45RB
CD2
CD44
CD38
ICAM-1
DNAM-1
CD49e
CD11c
E
Percentage
OR
non−OR
CR
PR
SD
PD
CR-SD: p=0.04
SD-PD: p=0.02
SD-PD: p=0.04
F
Up in OR
Up in non-OR
−log10(BH−adjusted p−value)
Effect Size
G
Up in OR
Up in non-OR
−log10(BH−adjusted p−value)
Effect Size
H
Up in OR
Up in non-OR
−log10(BH−adjusted p−value)
Effect Size
I
Up in OR
Up in non-OR
−log10(BH−adjusted p−value)
Effect Size

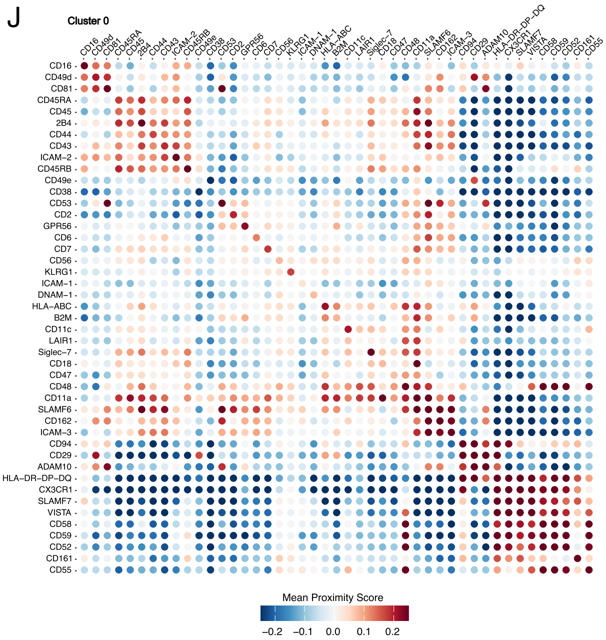


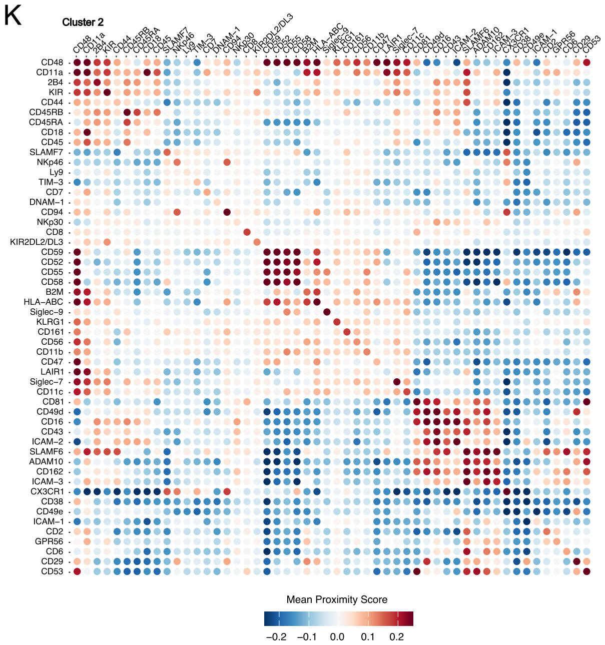


***Figure 5 Supplement. Differential analysis and composition of the NK cell compartment.*** *(**A**) UMAP projection of classic NK cell subpopulations. (**B**) Average relative frequency (%) of classic NK cell subpopulations within the total NK cell compartment. (**C**) Relative frequency (%) of classic NK cell subpopulations within the total NK cell compartment; split by outcome (top row) and clinical response (bottom row). P-values < 0.1 are indicated. (**D**) Heatmap of protein abundance, expressed as Z-scaled centered log-ratio (CLR) values, for the most variable proteins across identified subpopulations. (**E**) Relative frequency of the NK cell subpopulations detected by re-clustering (see Figure 5A) within the total NK cell compartment; stratified by outcome (top row) and clinical response (bottom row). P-values < 0.1 are indicated. (**F**) Volcano plot illustrating differential protein abundance between OR and non-OR within the whole NK cell compartment. (**G**) Volcano plot illustrating differential proximity scores between OR and non-OR within the whole NK cell compartment. Volcano plots illustrating differential abundance (**H**) and differential proximity scores (**I**) between OR and non-OR within the 3 largest NK cell clusters. Horizontal line denotes adjusted p-value < 0.05. Median per-patient mean proximity scores (log2 ratio) for relevant proteins within NK cell cluster 0 (J) and NK cluster 2 (K).*
